# Study of excess manganese stress response highlights the central role of manganese exporter Mnx for holding manganese homeostasis in the cyanobacterium *Synechocystis sp.* PCC 6803

**DOI:** 10.1101/2024.08.16.608223

**Authors:** Mara Reis, Sanja Zenker, Prisca Viehöver, Karsten Niehaus, Andrea Bräutigam, Marion Eisenhut

## Abstract

Cellular levels of the essential micronutrient manganese (Mn) need to be carefully balanced within narrow boarders. In cyanobacteria, sufficient Mn supply is critical for assuring the function of the oxygen-evolving complex as central part of the photosynthetic machinery. However, Mn accumulation is fatal for the cells. The reason for the observed cytotoxicity is unclear. To understand the causality behind Mn toxicity in cyanobacteria, we investigated the impact of excess Mn on physiology and global gene expression in the model organism *Synechocystis* sp. PCC 6803. We compared the response of the wild type and the knock-out mutant in the manganese exporter (Mnx), Δ*mnx,* which is disabled in the export of surplus Mn and thus functions as model for toxic Mn overaccumulation. While growth and pigment accumulation in Δ*mnx* was severely impaired 24 h after addition of 10-fold Mn, the wild type was not affected and thus mounted an adequate transcriptional response. RNA-seq data analysis revealed that the Mn stress transcriptomes were partly resembling an iron limitation transcriptome. However, the expression of iron limitation signature genes *isiABDC* was not affected by the Mn treatment, indicating that Mn excess is not accompanied by iron limitation in *Synechocystis.* We suggest that the Ferric uptake regulator, Fur, gets partially mismetallated under Mn excess conditions and thus interferes with an iron-dependent transcriptional response. To encounter mismetallation and other Mn-dependent problems on protein level, the cells invest into transcripts of ribosomes, proteases, and chaperones. In case of the Δ*mnx* mutant the consequences of the disability to export excess Mn from the cytosol manifest in additionally impaired energy metabolism and oxidative stress transcriptomes with fatal outcome. This study emphasizes the central importance of Mn homeostasis and the transporter Mnx’s role in restoring and holding it.

## INTRODUCTION

All organisms rely on an adequate manganese (Mn) supply to maintain the functions of enzymes, such as glycosyl transferases, oxalate oxidase, or Mn-dependent superoxide dismutase [1, 2]. Organisms performing oxygenic photosynthesis have in comparison to non-photosynthetic organisms a 100-fold higher demand for Mn since they use Mn for the oxidation of H_2_O at the <u>o</u>xygen-<u>e</u>volving <u>c</u>omplex (OEC) [3]. The OEC is a central part of <u>p</u>hoto<u>s</u>ystem <u>II</u> (PSII) and hosts the catalytic Mn cluster (Mn_4_CaO_5_), stabilized by PsbO, PsbP, and PsbQ in plants and PsbO, PsbU, and PsbV in cyanobacteria, respectively [4–8]. To ensure proper provision of Mn to the OEC, the model cyanobacterium *Synechocystis* sp. PCC 6803 (hereafter *Synechocystis*) maintains cellular Mn homeostasis. In a light dependent manner, Mn is imported via outer membrane pores with a low selectivity to the periplasm [4, 9]. Here, 75 % of the cell’s Mn pool are stored and bound either to the outer membrane or <u>Mn c</u>upin <u>A</u> (MncA) [4, 8]. The remaining 25 % are located in the cytoplasm or in the thylakoid system [3, 8, 10] and are associated with nucleic acids and small molecule chelates or bound to different metalloproteins [11]. To match the high Mn demand of the OEC, Mn is transported into the cytoplasm by two different Mn import systems. Recently, two members of the <u>u</u>nknown <u>p</u>rotein family 0016 (UPF0016), the <u>h</u>emi <u>m</u>anganese e<u>x</u>changers (Hmx) 1 and 2 were demonstrated to facilitate constitutive Mn uptake via the plasma membrane [12]. Upon limited Mn supply (< 1 µM, [13]), Hmx1/2 uptake is assisted by the high-affinity ABC-type transporter MntCAB (<u>Mn</u> transporter), which is transcriptionally regulated by the ManSR (<u>man</u>ganese <u>s</u>ensor / regulator) two-component system [13–15]. If Mn supply is sufficient (> 1 µM) Mn^2+^-ions bind to amino acid residues in the periplasmic loop of the sensor protein ManS. This leads to autophosphorylation of ManS, which phosphorylates the response regulator ManR subsequently. Phosphorylated ManR binds to the promoter of the *mntCAB* operon, blocking its expression. On the contrary, when Mn is scarce, ManS is not activated by phosphorylation, ManR not phosphorylated and the *mntCAB* operon is expressed, leading to increased import of Mn [13, 14]. Cytoplasmic Mn is either used in the cytoplasm by Mn-requiring enzymes or is further distributed to the thylakoid lumen by the <u>Mn</u> e<u>x</u>porter (Mnx) also known as *<u>Syn</u>echocystis* <u>P</u>hotosynthesis <u>A</u>ffected <u>M</u>utant 71 (SynPAM71), another member of the UPF0016 [10, 16]. Though it is questionable whether Mn limited conditions occur in aquatic environments, in lab experiments very low Mn supply (< 0.1 µM) leads to decreased photosynthetic activity since the H_2_O oxidation capacity of the OEC is lowered and as a consequence the overall growth rate is reduced [17].

In contrast to Mn limitation, Mn excess has not been studied in detail in cyanobacteria, yet. A surplus of Mn results in decreased chlorophyll *a* content and reduced <u>p</u>hoto<u>s</u>ystem I (PSI) activity on the physiological level and eventually leads to cell death in *Synechocystis* [10, 16]. The mechanism of this Mn toxicity is not well understood. Besides induction of iron (Fe) limitation, the most plausible mode of action is the mismetallation of enzymes and regulatory proteins, changing or abolishing their activity [8, 18]. According to the Irving-Williams series (Mg^2+^ < Mn^2+^ < Fe^2+^ < Co^2+^ < Ni^2+^ < Cu^2+^ > Zn^2+^), different metal ions compete with each other to be bound by amino acid residues [8, 19]. Correct metalation is only favored due to strictly controlled concentrations of the different metals at the site of metal incorporation during or after protein biosynthesis [20]. The vital importance for controlling the intracellular Mn concentration and subcellular allocation could be demonstrated for the mutant in the thylakoid Mn transporter Mnx [10]. Mnx transports Mn from the cytoplasm into the thylakoid lumen, where the OEC and the highest demand for Mn supply is located. The knock-out mutant Δ*mnx* displays high light sensitivity and a significantly longer recovery time compared to the <u>w</u>ild type (WT) after photoinhibition, presumably due to the lack of Mn in the thylakoid lumen for enabling the high D1 turnover [10]. It was shown that the Δ*mnx* mutant is highly sensitive towards Mn excess conditions in general and displays a lethal phenotype upon Mn stress as it accumulates Mn intracellularly [10]. Obviously, the subcellular Mn pools need to be carefully maintained at constant levels to ensure proper cell growth and Mnx plays a critical role in the correct subcellular Mn distribution [10]. However, it is not understood why the cytoplasmic Mn overload in *Synechocystis* is detrimental.

In this study, we aimed at a mechanistic understanding of the Mn excess response in the cyanobacterium *Synechocystis.* To this end, we grew *Synechocystis* cells under standard (1x) MnCl_2_ and excess (10x) MnCl_2_ conditions and investigated physiological and transcriptional effects. We studied the WT and the mutant Δ*mnx*, which is defective in the export of Mn from the cytosol. The WT survives 10x Mn and hence displays an adequate transcriptional response. Δ*mnx* succumbs to 10x Mn and shows a similar though more pronounced transcriptional response in comparison to the WT. According to our results, Mn excess induces a transcriptional Fe acclimation response with significantly reduced PSI and PSII, phycobilisomes, and Fe importer transcript abundances in both cell lines. We suggest that mismetallation of the transcriptional regulator <u>F</u>erric <u>u</u>ptake regulator (Fur) is one causative factor. In addition, the Δ*mnx* mutant displays a significant transcriptional reduction of ATPase and carbon metabolism genes in general and shows features of a response towards oxidative stress. Protective mechanisms are not sufficient to compensate for the Mn-dependent mismetallation, energy depletion, and reactive <u>o</u>xygen <u>s</u>pecies (ROS) generation in the Δ*mnx* mutant with fatal outcome.

## METHODS

### Cyanobacterial strains and growth conditions

A *Synechocystis* sp. PCC 6803 glucose-tolerant (Japan) strain served as the WT. The Δ*mnx* mutant line was generated in a previous study by insertion of a kanamycin resistance cassette into the *mnx (sll*0615) open reading frame [10]. Cells were grown in BG11 medium, pH 7.5 [21]. Cultivation in Erlenmeyer flasks was performed on a shaker at 100 rpm, 30 °C, and continuous LED illumination at an intensity of 100 µmol photons m^-2^ s^-1^. For the Δ*mnx* mutant line, the medium was supplemented with 50 µg µL^-1^ kanamycin. Before sampling for RNA isolation, cells were resuspended in fresh medium, adjusted to an optical density at 750 nm (OD_750_) of 0.3 and cell suspensions split into two flasks per strain. BG11 medium in one flask was supplemented with standard concentration of MnCl_2_ (1x Mn, 9 µM MnCl_2_). BG11 medium in the other flask was supplemented with excess MnCl_2_ (10x, 90 µM MnCl_2_). All treatments were performed in biological triplicates. After 24 h cultivation, 10 mL samples were taken from each culture and centrifuged in pre-cooled tubes for 10 min at 3,000 rpm, 4 °C. Cell pellets were snap-frozen in liquid N_2_ and stored at -80 °C until further use.

### Drop tests

For the drop test assays, *Synechocystis* cultures were grown as described above in BG11 medium with standard concentration of 9 µM MnCl_2_ and adjusted to an OD_750_ of 0.2. Dilutions of the cell suspensions 1:10, 1:100, and 1:1,000 were prepared and 2 µL of each dilution dropped onto BG11 agar plates [21] containing different concentrations of MnCl_2_ and/or Fe-NH_4_-citrate. Afterwards, plates were grown for 5 d at 30 °C under continuous illumination of 100 µmol photons m^-2^ s^-1^.

### Growth performance and pigment measurements

To determine the growth rate of WT and Δ*mnx* under Mn standard conditions and Mn excess conditions, growth curves were generated. For this approach, liquid cultures were grown as described above in Erlenmeyer flasks on a shaker. After precultivation, two 50 ml batches of WT and Δ*mnx* mutant with an OD_750_ of 0.2 were filled into growth tubes for the Multi-Cultivator MC-1000-OD (Drásov, Czech Republic, Photon Systems Instruments) and 9 µM or 90 µM MnCl_2_ were added to one tube of WT and Δ*mnx*, respectively. The cultures were grown for 7 d bubbled with filtered ambient air under constant illumination of 100 µmol photons m^-2^ s^-1^ at 30 °C. To estimate the growth rates, the OD_750_ was recorded every 20 minutes. The growth rate was determined during the logarithmic growth phase (0 h to 24 h and 24 h to 96 h) using the formula:

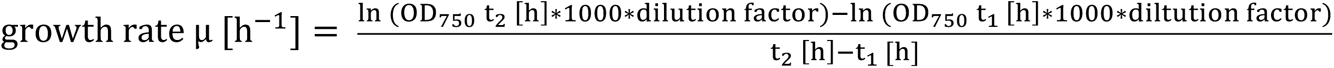

To calculate the chlorophyll *a*, phycocyanin, and carotenoid content, OD_680_ (chlorophyll *a*), OD_625_ (phycocyanin), and OD_490_ (carotenoids) were measured every 24 h in triplicates. Pigment contents were estimated according to [22].

### RNA-seq analysis

RNA was extracted from cell pellets using the Qiagen® RNeasy Plant Mini Kit (Hilden, Germany, Qiagen GmbH) following the manufactureŕs instructions with 10 µL 2-mercaptoethanol per 1 mL RLT buffer. For cell lysis, 500 µL beads with a size of 0.2 – 0.4 µm were used in a Precyllis Evolution cell lyser (Montigny-le-Bretonneux, France, Bertin Technologies) for 4 x 30 s at 2,000 rpm with 15 s pauses in between. RNA was prepared according to the Illumina® TruSeq Stranded Total RNA With Illumina Ribo-Zero Plus (Hayward, USA, Illumina). Procedure of rRNA depletion and preparation of the library were conducted as stated in the protocol of the corresponding reference guide from Illumina. The library-pool was sequenced using the Illumina® NextSeq500 with 76 BP SR (single read) at a HighOutput Flowcell.

All data analyses were performed with R 4.3.0. Differential gene expression analysis was performed with edgeR using the classic test followed by Benjamini-Hochberg multiple hypothesis correction [23, 24]. A principal component analysis was performed on transcript <u>p</u>er <u>m</u>illion (TPM) values using the prcomp function with parameters scale = T and center = T. Loadings were calculated, the six highest absolute values for component 1 and 2 were selected, and colored by the highest level KEGG <u>K</u>yoto <u>E</u>ncyclopedia of <u>G</u>enes and <u>G</u>enomes (KEGG) [25] map annotation.

### KEGG Enrichment

<u>K</u>EGG <u>O</u>ntology (KO)-term annotations were retrieved from eggNOG-mapper [26] with default parameters, KAAS [27] against *Synechocystis* sp. PCC 6803, *Synechocystis elongatus* PCC 7942, *Nostoc sp*. PCC 7120, and *Arabidopsis thaliana,* as well as directly downloaded from KEGG. The coding sequences from *Synechocystis* sp. PCC 6803 were used to obtain KO-term annotations from KAAS and eggNOG. Enrichments of significantly differentially abundant transcripts were tested on the “KEGG map” annotation level as well as the “KEGG module” level for the mutant and control, using Fisher’s exact tests [28]. Up-regulated genes were defined as those with a log_2_-fold change > 0 with *q*-value < 0.05 and down-regulated genes as log_2_-fold change < 0 and *q*-value < 0.05. Enrichments on the PCA eigenvalues were done as described above separately for positive and negative values using a cut-off of 0.015 and Fisher’s exact test. Enrichments with a *P*-value ≤ 0.05 were considered significant.

### Data availability

All code used in this analysis is available on GitLab (https://gitlab.ub.uni-bielefeld.de/computationalbiology/mn-excess-rna-seq; will be made public upon publication). The RNA-seq data set is available at the <u>E</u>uropean <u>N</u>ucleotide <u>A</u>rchive (ENA) with the project ID PRJEB75422. A table with transcript abundances and statistical data for all genes is provided in Table S1.

## RESULTS

### Mn excess impairs growth and pigment accumulation of Δ*mnx*

To assess the physiological impact of different Mn regimes on WT and the Δ*mnx* mutant line, growth rates and pigment contents were determined. To this end, cells were inoculated into BG11 medium supplemented with either the standard concentration of MnCl_2_ (1x Mn, 9 µM MnCl_2_) or with an excess concentration of MnCl_2_ (10x Mn, 90 µM MnCl_2_) and grown for 4 d. A pale green phenotype was observed for the Δ*mnx* mutant in comparison to the WT under both standard growth conditions and Mn stress conditions (Fig. 1A). The growth rates for the first 24 h after transfer to excess Mn medium did not significantly differ between all cultures (Fig. 1B). However, the growth rate determined for the 24 h-to-96 h interval demonstrated a significantly depleted growth of the Δ*mnx* mutant when grown under Mn excess (10x Mn) conditions (Fig. 1C). The rate was highly variable for the Δ*mnx* mutant in standard concentrations (Fig. 1C). The WT was not affected in growth by the elevated levels of Mn. With respect to pigment levels, the Δ*mnx* mutant grown in Mn excess accumulated significantly lower levels of chlorophyll *a* (Fig. 1D) and phycocyanin (Fig. 1E) over time. The carotenoid level was slightly but significantly reduced only after 48 h of growth under Mn excess conditions in the Δ*mnx* mutant (Fig. 1F). On the basis of these results we selected the timepoint 24 h after the experimental start to study effects of Mn treatment on the transcriptome of WT and mutant to avoid pleiotropic cytotoxic effects. At this time point Δ*mnx* shows first symptoms, such as reduced pigmentation, but still grows WT-like.

**Fig. 1:**
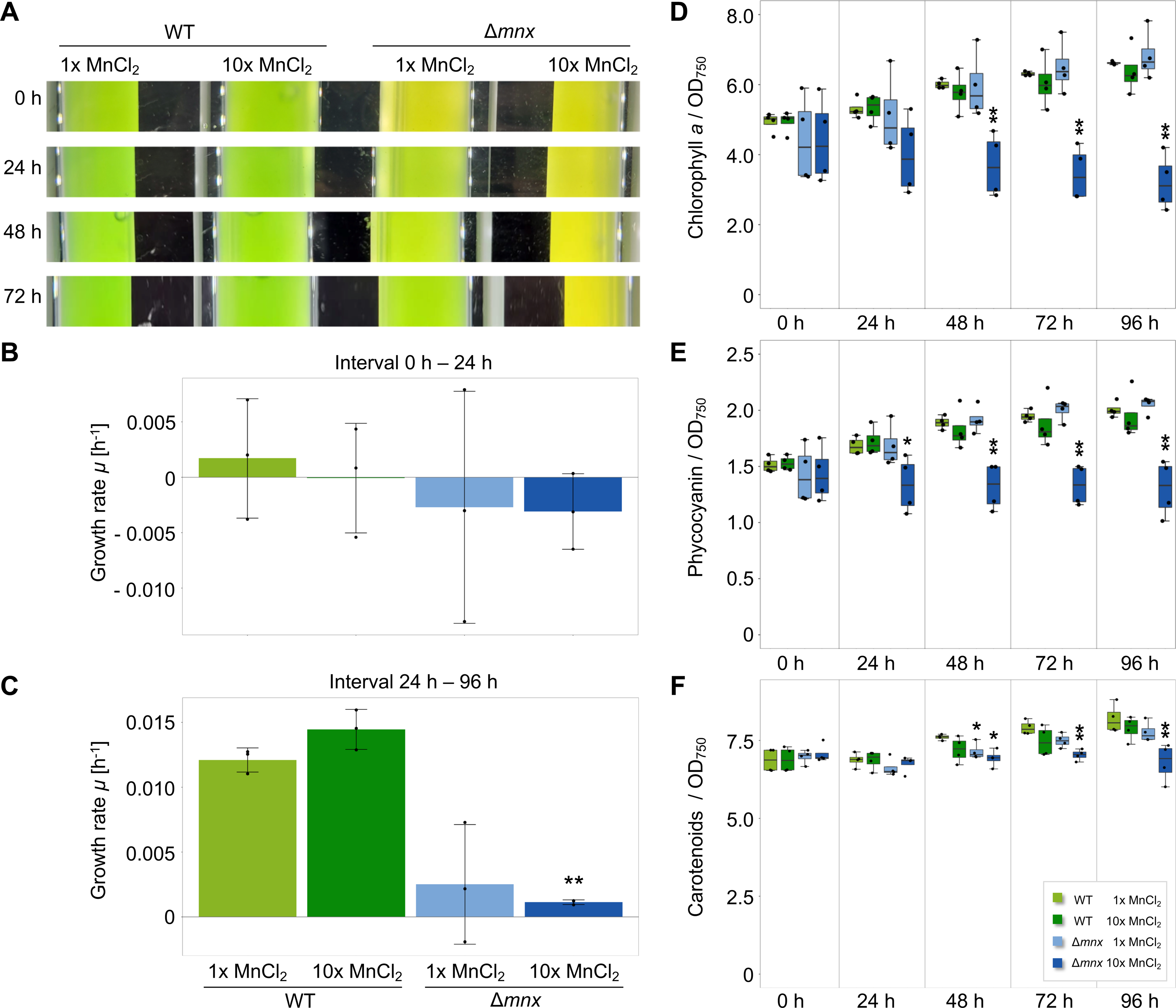
Effects of Mn treatment on growth and pigment content. **(A)** Phenotypic appearance of WT and Δ*mnx* mutant line under MnCl_2_ standard (1x, 9 µM MnCl_2_) and excess (10x, 90 µM MnCl_2_) conditions at different time points in the MC-1000-OD Multi-Cultivator. **(B), (C)** Growth rates of WT and Δ*mnx* mutant line under different MnCl_2_ regimes. Shown are growth rates for the first 24 h (0 – 24 h interval) (B) and the 24 h-to-96 h interval (C). **(D)** Chlorophyll *a* content, **(E)** Phycocyanin content, **(F)** Carotenoid content in WT and Δ*mnx* cells grown under different MnCl_2_ regimes. Pigment levels were normalized to the optical density at 750 nm (OD_750_). Significance in (B)-(F) was evaluated with Student’s t-Test *P* ≤ 0.05 (*); *P* ≤ 0.01 (**).

### Transcriptional profile of the Δ*mnx* mutant is more strongly affected by Mn excess than the WT

To assess the effect of excess Mn on the transcriptomes of WT and the Δ*mnx* mutant, three independent biological transcriptomes for each line were analyzed after 24 h growth under standard Mn and excess Mn treatment. The <u>p</u>rincipal <u>c</u>omponent <u>a</u>nalysis (PCA) of the transcript abundances does not show strong transcriptome differences between WT and mutant under 1x Mn treatment as their samples cluster together (Fig. 2A). The PCA revealed the Mn treatment as the major factor (PC1) contributing to 44 % of the variation, while the intracellular Mn allocation likely accounts for the 18 % variation in PC2 (Fig. 2A). A larger distance between the Mn excess treated Δ*mnx* to 1x samples compared to the WT replicates was observed in the first dimension. In addition, WT samples under excess Mn clustered away from all others in the second dimension. This result indicates a quantitative difference in the transcriptional response upon Mn excess in WT and Δ*mnx* cells and a qualitative difference in the response of WT to excess Mn. To gain first insight into the major contributing genes and their functional classification, component loading of the PCA was extracted and the first twelve components examined (Fig. 2B, Table S2). For PC1, *hemF* (*sll*1185, assigned to KEGG map “Porphyrin metabolism”), *cpgG* (*slr*2051, “Photosynthesis proteins”), a protein of unknown function (*slr*0554, “NA”), *apcC* (*ssr*3383, “Photosynthesis proteins”), *por* (*slr*0506, “Porphyrin metabolism”), and *psbA2* (*slr*1311, “Photosynthesis proteins”) have the strongest absolute impact. For PC2, *pstC1* (*sll*0681, “Transporters”) and *pstA1* (*sll*0682, “Transporters”), *feoB* (*slr*1277, “Transporters”), *pilQ* (*slr*1277, “Bacterial motility proteins”), and two proteins of unknown function (*sll*0182 and *sll*1106, “NA”) have the biggest absolute influence (Fig. 2B, Table S2). An enrichment of transcript’s eigenvalues shows enrichments for PC1 for “Photosynthesis”, “Carbon fixation, and “Ribosomes” and for PC2 for “Ribosomes”, “Photosynthesis, “Prokaryotic defense”, and “Transporters” (Table S3).

**Fig. 2:**
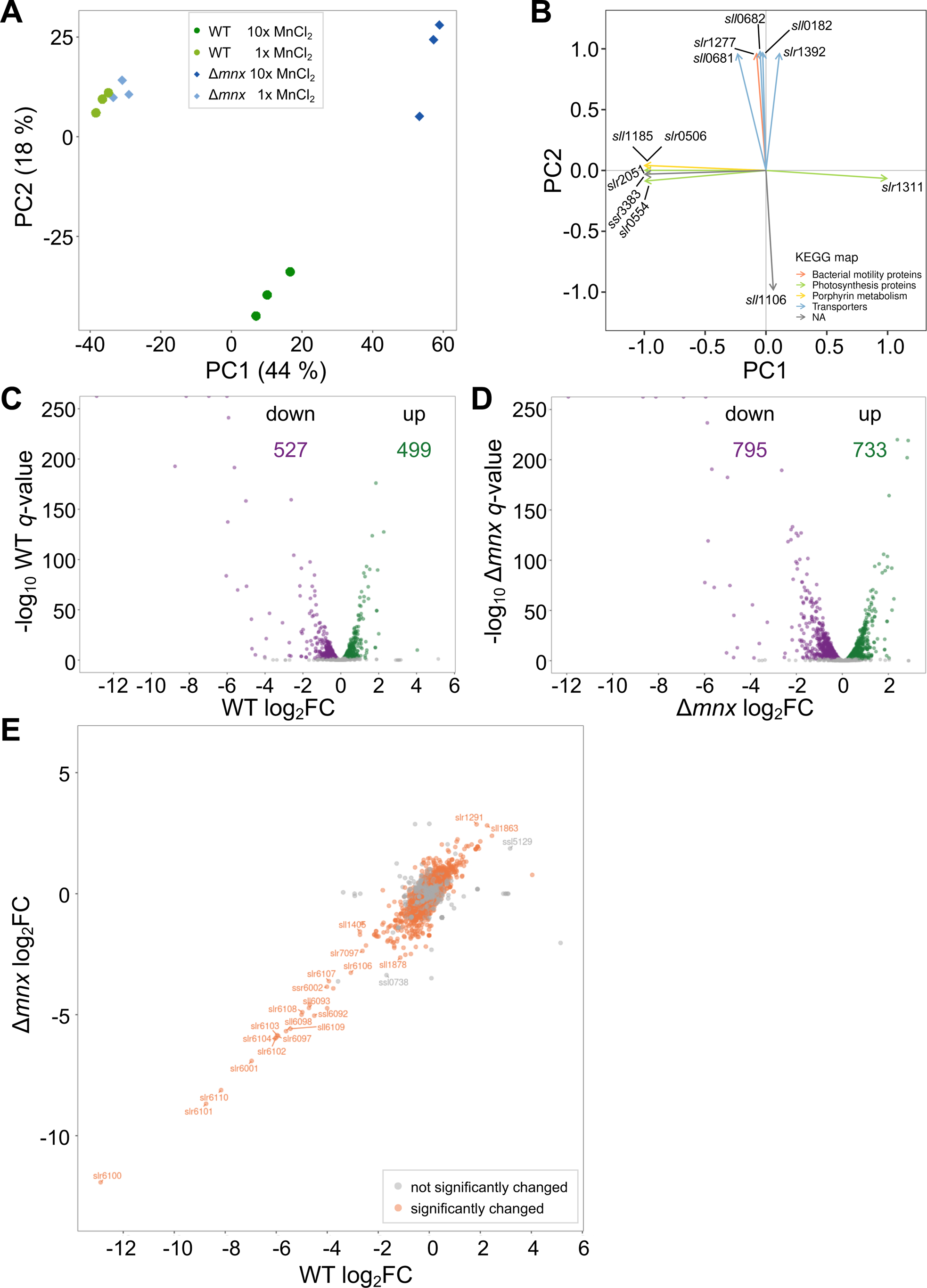
Effect of MnCl_2_ treatment on transcriptomes of WT and Δ*mnx*. **(A)** Principal component analysis (PCA) of transcript abundances in WT and Δ*mnx* cells grown under control and Mn excess (+Mn) conditions. **(B)** Component loading of the PCA. Eigenvectors of principal component 1 (PC1) and 2 (PC2) are shown with the corresponding gene loci. The color of the eigenvectors is according to the KEGG map of the respective gene, as displayed in the legend. Table S2 gives further information regarding the respective log_2_-fold changes of the gene and the assigned gene function. **(C)+(D)** Volcano plots of the global transcriptome responses of WT **(C)** and Δ*mnx* **(D)** towards different MnCl_2_ treatments. Shown are log_2_-fold changes (log_2_FC) of Mn excess (10x, 90 µM MnCl_2_) *versus* Mn control (1x, 9 µM MnCl_2_) conditions. Significant changes (*q* < 0.01; edgeR, [23]) are plotted in green (up) or violet (down) respectively. The number of significantly up- and downregulated genes is given for each genotype. **(E)** Scatterplot of the transcript abundances as log_2_FC in WT and Δ*mnx*. Orange color indicates significantly different (*q* ≤ 0.01) transcript abundances in WT and Δ*mnx* upon MnCl_2_ treatment, grey represents not significant differences. Gene loci of transcripts with a log_2_FC of ≥ |2.5| are displayed.

As suggested by the PCA, comparison of the number of <u>d</u>ifferentially <u>e</u>xpressed <u>g</u>enes (DEGs) upon Mn excess treatment to standard Mn concentrations showed differences between WT and mutant. In the WT, 1,026 transcripts were found significantly changed (*q* < 0.01, 27.8 % of all genes), with 499 genes showing enhanced transcript abundances and 527 genes showing reduced transcript abundances (Fig. 2C) 24 h after Mn excess treatment. Cells of the Δ*mnx* mutant had 1,528 significantly (*q* < 0.01, 41.4 % of all genes) changed transcripts, with 733 genes showing enhanced transcript abundances and 795 genes showing reduced transcript abundances (Fig. 2D). Calculating DEGs between genotypes within a treatment resulted in 25 significantly changed (*q* < 0.01, 0.7 % of all genes) genes under Mn control conditions (Fig. S1A, Table S4-1) and 862 significantly changed (*q* < 0.01, 23.3 % of all genes) genes under Mn excess conditions (Fig. S1B, Table S4-2), which agrees with the clustering of samples in the PCA (Fig. 2A). A scatterplot of the analyzed transcript abundance in WT and Δ*mnx* showed an apparently linear relationship between the transcriptional response of both WT and Δ*mnx* (Fig. 2E). The genes with the strongest transcriptional response had similar magnitude of response in mutant and WT while the genes with a more modest response between 4-fold up and down showed a stronger response in the mutant (Fig. 2E). The transcripts most impacted by Mn excess in both genotypes are mostly uncharacterized genes located on the extrachromosomal plasmids pSYSM, pSYSA and pSYSX (log_2_-fold change > |1.5|, Table S5. In addition, also *futC* (*sll*1878) and *exbD1* (*sll*1405), which are Fe responsive genes [29] were severely affected in their transcript abundance (Table S5). A small subset of transcripts falls away from the linear relationship (Fig. 2E) and likely represents those loading the PC2 of the PCA (Fig. 2A). These 81 transcripts include 20 with higher abundance in WT but lower in the mutants, such as *cbbL* (*slr*0009) or *petC* (*sll*1316), and 61 with significantly lower abundance in WT but higher in the mutant, such as *hliA* (*ssl*2542) or *ycf64* (*slr*1846) (Table S6). With the exception of a transposase and a gene without annotation, all transcripts are changed less than 2-fold in WT (Table S6).

These results indicate that WT and Δ*mnx* display a shared response to Mn stress, but the Δ*mnx* mutant line is more affected by Mn excess and the WT shows a small exclusive response.

### The functional transcriptome response largely overlaps between WT and Δ*mnx* mutant

For a detailed and mechanistic understanding of the Mn excess response, we analyzed the shared *versus* the strain specific transcriptional responses (DEGs with log_2_-fold change ≤ |1| and *q* ≤ 0.01) of WT and mutant.

WT and mutant share a response of 751 transcripts (Fig. 3A, B) despite displaying large differences in growth behavior (Fig. 1). The 400 shared less abundant transcripts represent 76 % of all less abundant WT transcripts but only 50 % of the genes with reduced transcript abundance in the Δ*mnx* mutant (Fig. 3A). Among the 351 shared transcripts with enhanced abundances, the overlap was similar with 70 % for the WT and 52 % for the Δ*mnx* mutant (Fig. 3B).

**Fig. 3:**
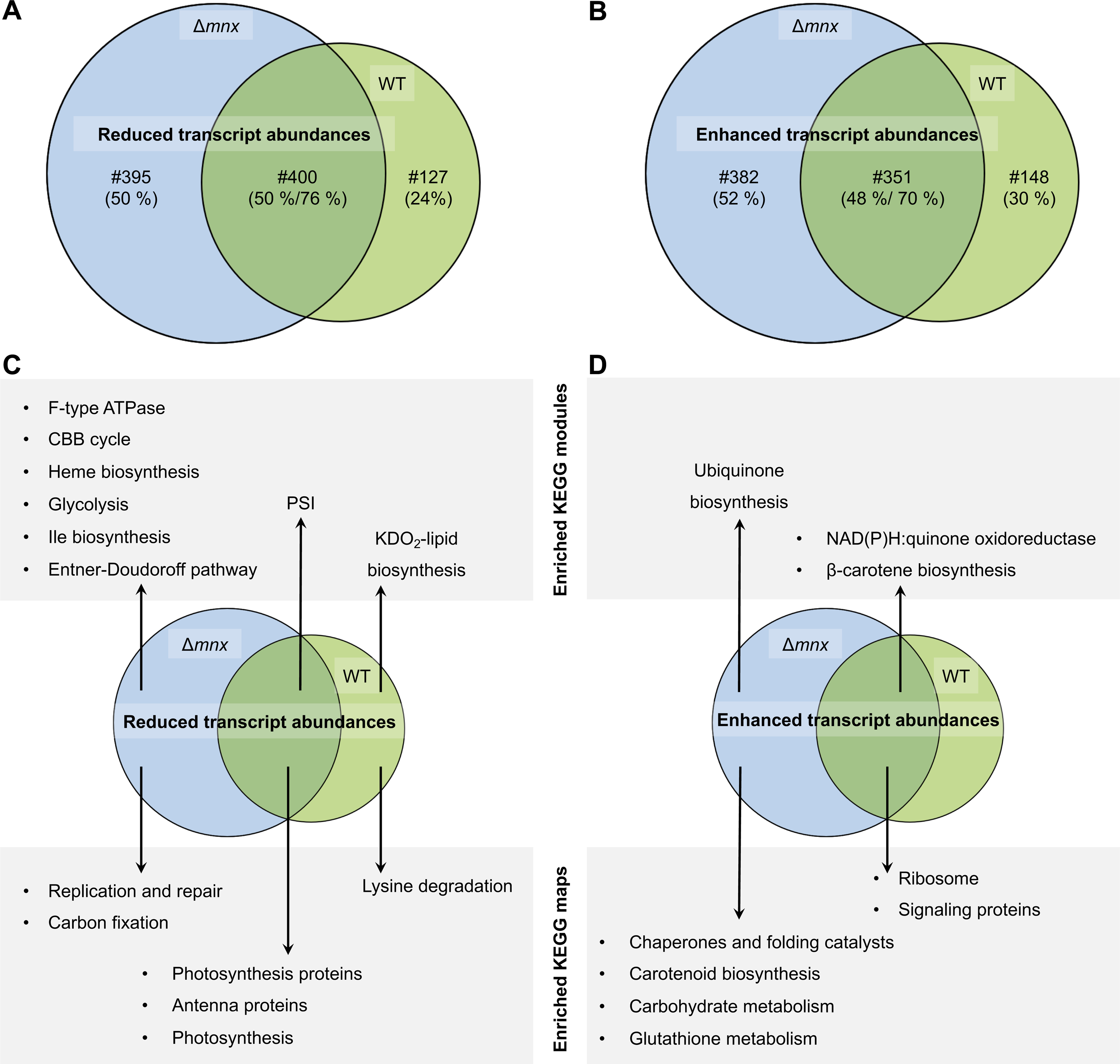
Comparison of the transcriptional response of WT and Δ*mnx* towards Mn excess. Overlap in transcriptional responses upon Mn excess (DEGs with log_2_-fold change ≤ |1| and *q* ≤ 0.01) is shown with Venn diagrams. The number (#) and percentage (%) of genes, which are shared or specific for Δ*mnx* and the WT, respectively, is given. **(A)** Overlap of genes with significantly reduced transcript abundances in WT and Δ*mnx* **(B)** Overlap of genes with significantly enhanced transcript abundances in WT and Δ*mnx*. **(C)** Overlap of significantly (*P* ≤ 0.05) enriched KEGG modules and maps in significantly reduced transcripts. **(D)** Overlap of significantly (*P* ≤ 0.05) enriched KEGG modules and maps in significantly enhanced transcripts.

To gain functional insights, we tested for enrichments of pathways using KEGG map annotations (Table S7) and sub-categories annotated as KEGG modules (Table S8). 24 h after Mn excess treatment, both WT and Δ*mnx* mutant showed for their genes with reduced transcript abundances (Fig. 3C) significant enrichment (*P* ≤ 0.05) of the KEGG maps “Photosynthesis”, “Photosynthesis proteins” and “Antenna proteins”. On the more specific KEGG module level, “PSI” was enriched for both strains. The biosynthesis of 3-deoxy-D-manno-octulosonic acid-lipid A (KDO_2_-lipid A), a component of the lipopolysaccharide layer of the outer membrane in gram-negative bacteria [30], was the only pathway that was downregulated in the WT according to KEGG modules enrichment as was lysine degradation. The Δ*mnx* mutant line solely showed enrichments of several pathways in C metabolism. The KEGG module enrichments (*P* ≤ 0.05) revealed reduced transcripts (Fig. 3C) corresponding to the modules “<u>C</u>alvin-<u>B</u>enson-<u>B</u>assham (CBB) cycle”, “Glycolysis”, “Entner-Doudoroff pathway”, and “F-type ATPase”, and corresponding to the maps “Ile biosynthesis”, “Heme biosynthesis”, “CO_2_ fixation”, “Replication and repair” only in the Δ*mnx* mutant line.

With regard to genes with enhanced transcript abundances upon Mn excess treatment (Fig. 3D), WT and Δ*mnx* mutant shared enrichment of the categories “NAD(P)H:quinone oxidoreductase”, “β-carotene biosynthesis”, “Ribosomes”, and “Signaling proteins”. While the specific WT response of 148 transcripts did not show any further enrichment, the Δ*mnx* mutant was affected additionally in “Ubiquinone biosynthesis”, “Chaperones and folding catalysts”, “Carotenoid biosynthesis”, “Carbohydrate metabolism”, and “Glutathione metabolism”.

### Mn importer MntCAB is reduced in transcript abundance upon Mn excess treatment

We hypothesized that Mn excess affects the expression patterns of genes that encode proteins involved in Mn homeostasis (Fig. 4, [8]). The gene transcripts of the low-Mn inducible high-affinity Mn importer MntCAB [31] were significantly less abundant in both WT and mutant. The second, constitutive Mn import system Hmx1/2 [12], was not significantly changed in transcript abundances in both strains. In the WT, also the Mn exporter gene *mnx* was not affected on transcript level. Transcripts of PratA, which is postulated to function in loading of the pre-D1 protein with Mn, was significantly more abundant with even higher induction in the Δ*mnx* mutant line [32]. In accordance, levels of *psbA2* and *psbA3*, encoding D1, were significantly upregulated in both genotypes but even higher in Δ*mnx* upon Mn excess treatment. For the Mn responsive two-component system ManSR, transcript abundances of ManS were unaffected, while transcript levels of ManR were significantly induced in the mutant only. Since metal transporters may be co-regulated under metal stress situations, we investigated also expression of metal ion transport systems for Ca, Cu, Mg, Mo, Ni, Co, Zi, and K (Table S9) [33]. We found for the genes *copBAC* significantly higher abundances in both cell lines. Copper resistance protein BAC (CopBAC) is a cation efflux transporter, which is known to export copper [33].

**Fig. 4:**
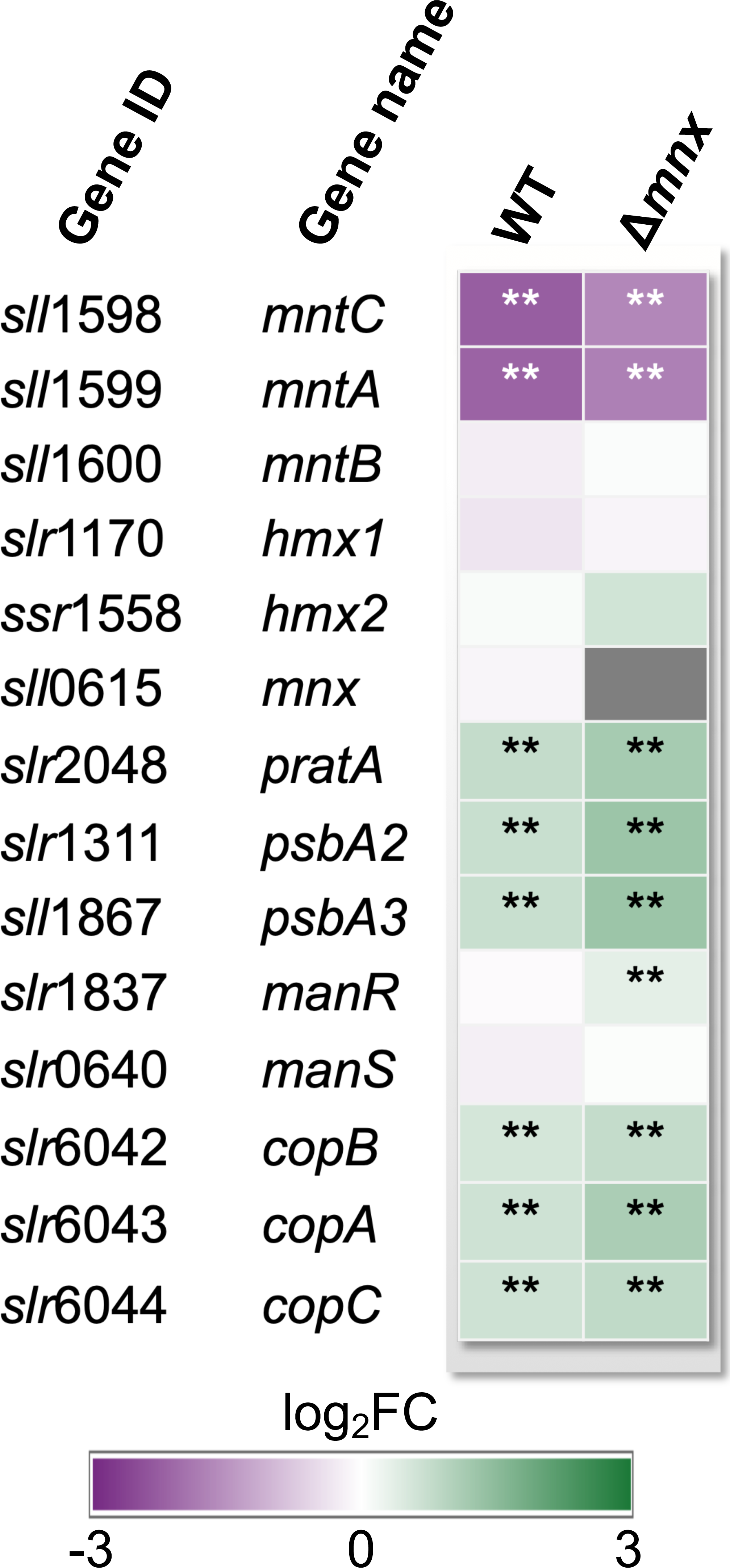
Transcriptional response of genes involved in Mn homeostasis. Transcriptional response upon Mn excess treatment is presented as heatmap of corresponding log_2_-fold change (log_2_FC) for the WT and Δ*mnx* mutant line. Statistical differences were evaluated according to Benjamini-Hochberg with *q* ≤ 0.05 (*); *q* ≤ 0.01 (**). Gene loci and names were obtained from [64]

### Transcriptional Mn excess response is partially congruent with Fe acclimation response

Since studies on *Escherichia coli* (*E. coli*) have demonstrated that Mn excess induces Fe limitation [18] and the most affected genes in our work are known to be Fe responsive (Fig. 2E), we hypothesized that the Mn excess transcriptomes show features of an Fe limitation response. To test this hypothesis, we compared our transcriptome data to transcriptomic data of Fe limitation in *Synechocystis* for major Fe-responsive pathways (Fig. 5, [29]). Transcript abundances of photosynthesis (phycobilisome, PSI, PSII, and ATPase) and CBB cycle corresponding genes were significantly reduced upon Mn treatment as they are in Fe limitation (Fig. 5). Three genes behaved opposite in Mn excess compared to Fe limitation, *psaA* was upregulated in Fe limitation but reduced in Mn excess, and for the D1 encoding genes *psbA2* and *psbA3* no information is available in Fe limitation but they were upregulated in Mn excess. Fe limitation induces chaperones and proteases [29, 34] and in Mn excess transcripts encoding chaperones and proteases were more abundant (Fig. 5). Other genes from known Fe limitation response pathways reacted under Mn excess conditions in a reverse manner. Transcripts encoding carboxysomal proteins, as part of the <u>c</u>arbon <u>c</u>oncentrating <u>m</u>echanism (CCM) had enhanced transcript levels under Mn excess (Fig. 5). Transcripts encoding the major Fe importer in the plasma membrane, ATP-binding cassette-type Fe(III) transporter FutABC, *futABC* were significantly less abundant under Mn excess conditions, while Fe limitation leads to enhanced transcript levels [35]. Transcript accumulation of other genes encoding Fe importers, such as <u>fe</u>rrous ir<u>o</u>n transport protein <u>B</u>, *feoB*, was significantly reduced under Mn excess in WT only. The energy transmitter genes *tonB* and *exbD1/B1* were significantly reduced in transcript abundance, compared to the control as were the genes for <u>o</u>uter <u>m</u>embrane channel <u>p</u>roteins (OMPs). The typical Fe limitation indicative genes iron <u>s</u>tress induced (isi) *isiA* and *isiB* were not significantly altered in their transcript abundances. The inconsistent pattern led to a non-significant overlap between the transcriptome responses of Mn excess and Fe limitation as tested by hypergeometric distribution calculation (Table S10).

**Fig. 5:**
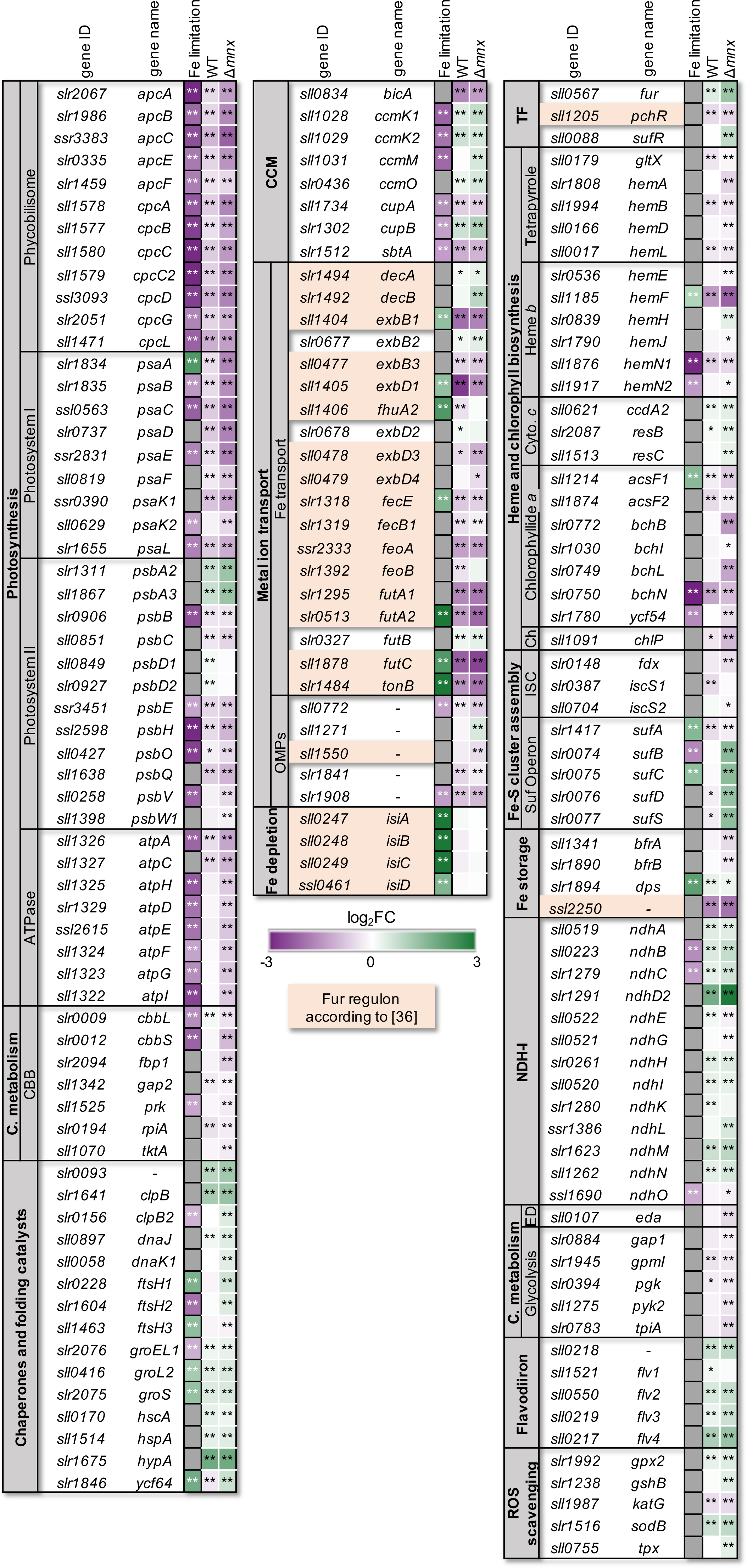
Comparison of Mn excess transcriptional response in WT and Δ*mnx* with Fe limitation response according to Singh et al. (2003) [29] in WT. Color coding of the boxes indicates the reduced (violet) or increased (green) transcript abundance of the corresponding gene(s) represented by the mean of the log_2_-fold changes (log_2_FC). Asterisks indicate significant changes with *: q-value < 0.05, **: q-value < 0.01. Values are given in Table S10. Some data points of the Fe limitation set exceed the color coded log_2_-fold change of |3| and are presented in deep violet and green for better comparison. Abbreviations: C.: carbohydrate, CBB: Calvin-Benson-Bassham cycle, CCM: carbon concentration mechanism, Ch: chlorophyll *a* biosynthesis, Cyto. c: cytochrome *c* biosynthesis, ED: Entner-Doudoroff pathway, Fe: iron, OMPs: outer membrane channel proteins, ROS: reactive oxygen species.

To get a fuller picture, we also investigated functional categories associated with above mentioned pathways but not specifically mentioned in the iron limitation data set [29]. Studying expression of transcription factors that are known to being intrinsic regulators of Fe homeostasis in *Synechocystis*, that are Fur, SufR, and PchR [36, 37] we found for *fur* and *sufR* significantly stronger transcript levels especially in Δ*mnx*, while *pchR* was reduced (Fig. 5). Connected to the light reactions of photosynthesis, we detected mostly reduced transcript levels of genes involved in heme and chlorophyll biosynthesis with the exception of cytochrome *c* biogenesis in Mn excess and mixed pattern in significantly changed transcripts the Fe limitation data (Fig. 5). Fe-S cluster biogenesis is an exception to shared WT and mutant excess Mn responses. It is majorly operated by the <u>s</u>ulfur <u>u</u>tilization factor (Suf) system in *Synechocystis* and was only enhanced on transcript level of the *sufBCDS* operon in the Δ*mnx* mutant again with mixed responses observed in Fe limitation (Fig. 5). With regard to Fe storage the transcriptional response to Mn excess is varied. The two <u>b</u>acterioferritin (bfr) family protein genes *bfrA* and *bfrB* [38] *w*ere both reduced on transcript level in Δ*mnx*. The bfr-associated ferredoxin encoding gene *ssl*2250 [36] was strongly down-regulated in both genotypes, while unaffected under Fe limitation. The gene *slr*1894, encoding an Fe storage protein of the Dps family [39], was enhanced in transcript abundance under both Fe limitation and Mn excess. The transcripts encoding subunits of <u>N</u>A<u>D</u>(P)<u>H</u>:chinon-oxidoreductase (NDH-1) complexes, which are central components of respiration, cyclic electron flow, and intracellular CO_2_ accumulation [40], were up-regulated in Mn excess with the sole exception of *ndhO* but opposite in Fe limitation for the two significant data points. Glycolytic enzymes were reduced in transcripts upon Mn excess treatment with no significant changes in Fe limitation. With regard to prevention of ROS formation, we observed enhanced transcript abundances of the <u>fl</u>a<u>v</u>odiiron proteins Flv2, Flv3, and Flv4, which serve as alternative electron sinks at the acceptor side of PSI [41] and no significant changes in Fe limitation. Furthermore, ROS scavenging by <u>s</u>uper<u>o</u>xide <u>d</u>ismutase (SOD) and glutathione appeared stronger on transcript level and again no significant changes in Fe limitation were determined.

Taken together, the comparison of the Mn excess transcriptome with Fe limitation transcriptome showed a partial overlap however with contrasting patterns to some extent, while the Fe-limitation signature genes *isiABCD* are unaffected under Mn excess conditions. Effects were stronger in the Δ*mnx* mutant.

### Fe surplus does not rescue the Mn excess phenotype in Δ*mnx*

Based on the results from the transcriptome data, which was in parts similar to a Fe limitation response, we hypothesized that additional Fe supplementation shall rescue growth of *Δmnx* upon Mn excess. Accordingly, we performed growth tests (Fig. 6) on BG11 plates with standard (1x Mn) or excess Mn concentrations (10x Mn) and increasing Fe-NH_4_-citrate concentrations (1x, 6 µg mL^-1^ Fe-NH_4_-citrate to 20x, 120 µg mL^-1^ Fe-NH_4_-citrate). While a growth difference between WT and *Δmnx* mutant at standard Mn concentration was not obvious, the lethal phenotype of the *Δmnx mutant* under Mn excess conditions could not be compensated by extra supplementation with Fe up to 20x (Fig. 6).

**Fig. 6:**
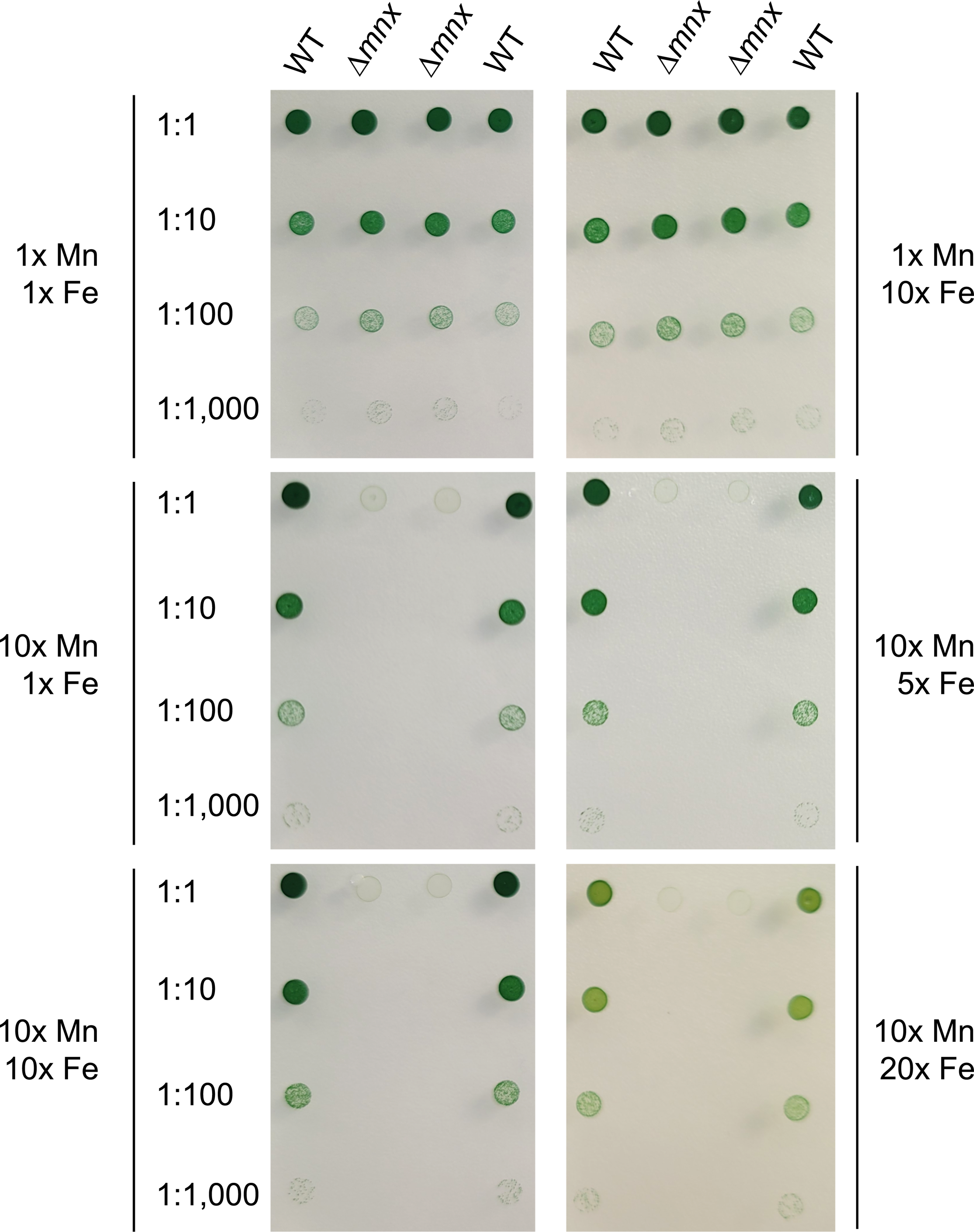
Growth of WT and Δ*mnx* mutant line under different Mn/Fe treatments. Growth of different dilutions (1:1, 1:10, 1:100, 1:1,000) were investigated on BG11 medium supplemented with standard Mn/Fe concentrations (1x Mn, 9 µM MnCl_2_; 1x Fe, 6 µg mL^-1^ Fe-NH_4_-citrate), standard 1x Mn and surplus Fe (10x Fe, 6 µg mL^-1^ Fe-NH_4_-citrate), or on excess Mn (10x Mn) and increasing Fe concentrations (1x, 5x, 10x, 20x Fe). Plates were photographed after 5 d incubation under continous illumination with 100 µmol photons m^-2^ s^-1^ at 30 °C.

## DISCUSSION

Mn toxicity is a poorly understood process. It is clear that the cellular Mn load needs to be controlled within narrow boarders. For single-cell organisms, such as *Vibrio cholerae*, *E. coli*, or *Synechocystis*, it was observed that already a 2- to 3-fold increased Mn-loading was lethal when the main Mn export system was knocked out [10, 16, 18, 42, 43]. So far, only for *E. coli* a detailed study of the Mn excess response has been performed [18]. In this organism Mn excess leads to Fe deficiency. As a consequence, Fe-S cluster assembly and heme biogenesis are impaired. This leads to a disruption of Fe-dependent electron transport chains and a block of the tricarboxylic acid cycle, causing an ATP crisis, which affects vital cellular processes. Additionally, the production of ROS induces DNA damage and affects protein stability [18]. In contrast to heterotrophic bacteria like *E. coli*, cyanobacteria have an at least 100-fold higher demand for Mn, since they utilize the micronutrient as the inorganic catalyst of light-driven water oxidation [4]. Thus, it is possible that Mn homeostasis and its regulatory network works differently.

### WT displays an adequate transcriptional response that reflects coping with Mn stress

The physiological response of the WT with stabile growth and pigmentation (Fig. 1) indicated that the WT was able to handle the extra Mn and took measures to restore Mn homeostasis after application of excess Mn and/or to function in excess Mn. Likely, the Mn exporter Mnx enabled efficient efflux of Mn surplus and recovery of at least adequate cellular Mn pools that, if combined with the observed transcriptional changes, allowed unaffected growth. According to our results, the transport capacity of Mnx is not regulated on the transcriptional level (Fig. 4). It is rather likely that posttranslational regulation might occur to regulate the activity of Mnx. Coping with the Mn stress implicates that the transcriptional profile we detected for the WT is an adequate response. The transcriptional response and anticipated consequences in *Synechocystis* are summarized in Fig. 7 and discussed in the following.

**Figure 7:**
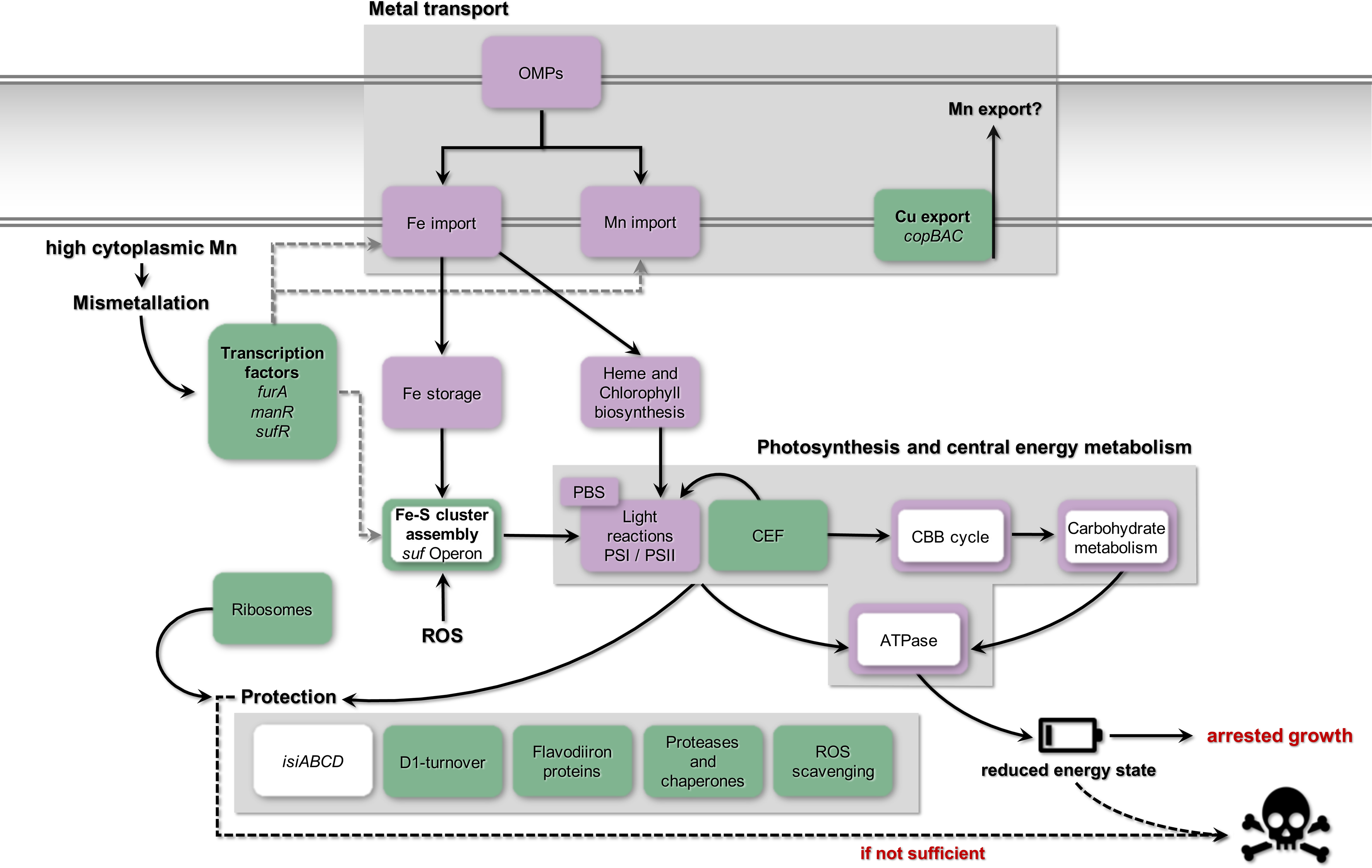
Model for a Mn excess response in *Synechocystis*. Presented are the effects of a Mn excess treatment on the global transcriptome of *Synechocystis* WT and Δ*mnx* mutant, which is defective in Mn efflux. The tile color indicates whether genes of the functional category were enhanced (green), reduced (violet), or unchanged (white) in transcript abundance 24 h after application of Mn excess stress in both, WT and Δ*mnx* mutant. Responses that are exclusive to the Δ*mnx* mutant are displayed by a color-framed tile. The response of specific transcripts is given in Fig. 5 and Table S1. Transcription factors related to Fe (Fur) and Mn import (ManR), and Fe-S cluster assembly (SufR) are transcriptionally up-regulated. In dependence of the cytoplasmic Mn status, Fur likely gets mismetallated with Mn instead of Fe and conveys a transcriptional response that is partially overlapping with an Fe acclimation response in *Synechocystis*. Cellular entrance of both Mn and Fe gets reduced due to down-regulation of Fe and Mn import systems as well as OMPs. Importantly, transcripts of the Fe deficiency responsive operon *isiABCD* are not altered, indicating that cells are not suffering from Fe limitation. Cu export via CopABC is upregulated on transcript level. We suggest that CopABC possibly assists Mn export in the Δ*mnx* mutant. However, this is a speculation and thus indicated with a question mark (?). Fe-dependent heme and chlorophyll biosynthesis is down-regulated on transcript level, overall leading to a lower abundance of photosynthesis, that is light-harvesting via phycobilisomes (PBS), light reactions and carbon reactions (CBB cycle), and carbohydrate metabolism related gene transcripts. Together with a downregulation of ATPase corresponding gene transcripts, cellular energy levels become depleted, with arrested cell growth as outcome. To cope with the detrimental effects of Mn excess, several protection mechanisms (D1 turnover, flavodiiron proteins, ROS scavenging, proteases and chaperons) are enhanced on transcript level to prevent cell death. In the case of the Δ*mnx* mutant the Mn exporter is not operative. The mutant is not able to adjust cytoplasmic Mn homeostasis. Consistently exclusive to Δ*mnx* mutant cells is the enhanced expression of the *sufBCDS* operon, involved in Fe-S cluster assembly, which indicates ROS stress in those cells upon Mn stress treatment. The protection mechanisms to deal with ROS and also mismetallation effects are insufficient and lead to cell death. Abbreviations: CBB cycle: Calvin-Benson-Bassham cycle, CEF: cyclic electron flow, CopBAC: copper resistance protein BAC, Cu: copper, Fe: iron, Fe-S: iron-sulfur cluster, Fur: ferric uptake regulator, ManR: Mn regulator, isi: iron stress induced, Mn: manganese, OMP: outer membrane channel protein, PBS: phycobilisome, PSI/II: photosystem I/II, ROS: reactive oxygen species, SufR: sulfur utilization factor regulator.

### Excess Mn treatment does not induce Fe limitation in *Synechocystis*

In general, the WT transcriptomes were affected by the Mn treatment after 24 h, though to a rather mild extent in comparison to the Δ*mnx* mutant. In accordance with Mn stress induced Fe depletion observed in *E. coli* [18] we detected features of a typical Fe limitation response in the cyanobacterium *Synechocystis.* This response included reduced transcript abundances of genes involved in chlorophyll and heme biosynthesis, photosynthesis covering both light and carbon reactions, carbon catabolism and respiration (Fig. 5, Fig. 7). However, the overlap between the Mn excess transcriptomes and a representative Fe limitation transcriptome [29] was not significant. We also compared our data with further Fe limitation transcriptomes [36, 44, 45] coming to the same result (Table S10). Not being Fe-limited is additionally supported by the unaffected levels of *isiABCD* (Fig. 5). Induction of this operon is considered as hallmark for Fe limitation in cyanobacteria [46]. We did not observe limitation in intracellular Fe content 4 h after high Mn treatment in an earlier experiment [10]. Thus, we suggest that the partially congruent response is not caused by the cellular Fe status but due to cross regulation of metal-responsive transcriptional regulators.

### Mismetallation of the transcriptional regulator Fur likely triggers parts of transcriptional response towards Mn excess

A reasonable explanation for the partially Fe-dependent transcriptional response is possible crosstalk of transcriptional regulators. Fur is generally considered as a key regulator in bacterial Fe homeostasis [47]. According to our data, the transcript abundance of *fur* is significantly increased in both cell lines (Fig. 5, Fig. 7), which was also described by [44] upon Fe limitation in *Synechocystis*. Many targets of the Fur regulon are affected during Mn excess conditions (Table S11), however in an unexpected way. The Fur-regulated Fe importers FhuA, FutABC, FeoAB, TonB/ExbBD, and FecBCDE are under Fe limitation typically up-regulated [36] but under Mn stress down-regulated (Fig. 5). The transcriptome data suggests that Fur itself commits besides an Fe-also a Mn-dependent response. In case of Fe-binding by Fur, the binding of one Fe^2+^-ion per monomer induces the dimerization of Fur and thereby enables the binding of the transcription factor to Fur boxes. Mostly, Fur acts as a repressor and *e.g*., represses expression of Fe importers upon Fe sufficiency, but can also act as indirect activator via the repression of regulatory asRNAs [47]. However, Fur was demonstrated to bind not only Fe but also Mn in *Bacillus subtilis*, that is mismetallation of Fur, leading to inappropriate repression of Fe uptake proteins [48]. This is in line with our observation of transcriptional reduction of Fe import proteins (Fig. 5, Fig. 7) and at least partially explains the transcriptional response being similar to an Fe acclimation response. It is furthermore conceivable that Mn-binding not only changes the activity of the transcriptional regulator with regard to repression/activation but also enables control of regulons different from those typically known for Fe-Fur. Mismetallation appears in the case of Fur to be rather advantageous and not a collateral damage.

Fur is also central to the regulation of the *isiABCD* operon. Typically, Fe limitation results in Fur-dependent de-repression of the *isiABCD* operon [36]. Thus, the unaffected transcript level of *isiABCD* support mismetallation of Fur as likely regulatory mechanism but also additional levels of regulation need to be considered. Transcript accumulation of *isiABCD* is regulated via Fur and the antisense RNA *iscR*, which is transcribed from the noncoding *isiA* DNA strand. Both mechanisms lead to a repression of either the *isiABCD* operon directly (Fur) or lead to a degradation of the corresponding RNA (*iscR*) [49]. Furthermore, the *isiABCD* operon is positively regulated via the oxidative stress responsive regulator RpaB [50] and negatively via the Fe regulator PfsR [51]. The gene transcripts of both transcriptional regulators were not altered in our transcriptome data (Table S1).

With regard to Mn-dependent transcriptional regulation heterotrophic bacteria mainly use two different mechanisms: Mn-binding transcription factors, i.e. MntR, and Mn-binding riboswitches, i.e. the *yybP-ykoY* riboswitch (reviewed in [11]). In contrast, little is known for cyanobacteria. So far, only the ManSR two-component system has been identified to being involved in regulation of Mn homeostasis in cyanobacteria [13–15, 44]. In agreement with the high external Mn concentration in our study, expression of the response regulator ManR was increased (Fig. 4), and the response regulator likely phosphorylated by ManS. As a result, binding of phosphorylated ManR repressed transcription of its primary target, the *mntCAB* operon, hampering further Mn uptake via this system (Fig. 4, Fig. 7).

Consistently with studies on Fe and/or Mn limitation [44], we observed cross-regulation of genes playing central roles in Mn and Fe homeostasis. We suggest a likely interconnection of Fur and ManR or yet unknown Mn-dependent regulators to integrate signals and orchestrate an appropriate transcriptional response enabling to finally deal with the stress situation. Future studies will reveal the function of transcriptional regulators in maintaining Mn homeostasis.

### Diminishment of Mn uptake comes along with reduced Fe uptake

According to [18], Mn stress in *E. coli* induces down-regulation on transcriptional level of Fe import and biosynthesis genes for the Fe siderophore enterobactin, leading to an Fe limitation phenotype. Upon extra addition of Fe to the medium, the transcript abundances of Fe import systems were raised and thus, it was possible to rescue the Mn stress phenotype in the *E. coli* mutant in the Mn efflux pump MntP [18]. We observed likewise downregulation of Fe import systems but were not able to compensate Mn toxicity in the *Synechocystis* Δ*mnx* mutant by supplementation with extra Fe (Fig. 6). We suggest that this observation is due to the occurrence of different uptake mechanisms in both organisms:

*E. coli* is an Fe-centric bacterium, which does not rely on Mn except for ROS scavenging. Hence, for Fe uptake several transporters exist, such as FecABCDE, FepBCDG, FhuA and FeoAB [52–54], while Mn import is facilitated by the highly specific Mn importer MntH [55]. Upon Mn limitation and oxidative stress, MntH expression is induced to facilitate Mn influx [56]. In contrast, cyanobacteria, such as *Synechocystis*, are dependent on both, abundant Fe and Mn supplies. Efficient Mn uptake to ensure Mn-dependent photosynthetic water-splitting activity [4] is realized by the use and interplay of the inducible high-affinity MntCAB and the constitutive Hmx1/2 [12] system at the plasma membrane and Mnx [10, 16] at the thylakoid membrane [57]. Also, the mainly Fe-transporting FutABC system likely supports Mn import, however in a low-affinity manner [8, 12, 44]. The entrance via the outer membrane into the periplasm is suggested to be shared between Fe, Mn, and other metals. According to the transcriptional profiles, the overaccumulation of Mn leads to decreased transcription of genes encoding the Mn import systems MntCAB (Fig. 4) and FutABC (Fig 5). As a result, further efficient Mn uptake is likely reduced or stopped to prevent the cell from damage due to the accumulation of intracellular Mn (Fig. 7). According to its suggested house-keeping function [12], transcript levels of the Hmx1/2 Mn transporter remained unaltered. The Mn exporter Mnx was not affected in the WT on transcriptional level (Fig. 4). Possibly, to fully abolish Mn uptake, also components for the uptake via the outer membrane, *tonB*, *exbB1, and exbD1* [36] were significantly lowered in transcript levels. The transcriptional repression is again explainable with mismetallated Fur, since it acts as transcriptional regulator of those genes. As a consequence, Fe uptake using the same outer membrane passage was hindered, too. Interestingly, Sharon and coworkers [44] already postulated a common transcriptional response of certain Fe and Mn transporters under Fe- and/or Mn-limiting conditions. A shared path of Fe and Mn was furthermore supported by the finding that addition of surplus Fe did not rescue the Δ*mnx* mutant from death under Mn excess conditions (Fig. 6). Furthermore, this result also fosters the notion that Fe limitation was not the reason for cell death of the Δ*mnx* mutant upon excess Mn treatment. The Fe uptake systems are down-regulated but not fully repressed on transcriptional level. Thus, extra Fe supply would have enabled enhanced Fe uptake by the cells and compensated a possible limitation phenotype. However, the treatment did not rescue the Δ*mnx* mutant and again supports together with unaffected expression of *isiABCD* (Fig. 5) that Fe limitation was not causing the fatal outcome of Mn excess treatment in the Δ*mnx* mutant.

The enhanced expression of *copBAC* (Fig. 4), which encodes a typically Cu-exporting transporter [33] is either explained by the co-regulation with Fe [58] or a hypothetical assignment of CopBAC as (low-affinity) Mn exporter. However, involvement in Mn transport remains to be tested.

### Detrimental effects of Mn excess are fought on several levels

Mismetallation is a common event when metal homeostasis is disturbed [59]. Prime targets for mismetallation are Fe-containing proteins. Hence is PSI, which is also transcriptionally down-regulated (Fig. 3C, Fig. 5), the major target of Mn excess within the PS apparatus of plants, such as *Arabidopsis thaliana* ([60]), and possibly also in *Synechocystis.* As a consequence, photosynthetic electron transfer is impaired and entails a highly delicate challenge for oxygenic photosynthetic organism. To prevent or deal with the formation of ROS, the transcriptomes inform about possible strategies *Synechocystis* employs (Fig. 7): i) Transcript levels of Flv proteins Flv2/3/4 are enhanced (Fig 5) to serve as an alternative electron sink at PSI [41]; ii) Transcript abundances of ROS scavenger proteins (SodB, GPX cycle, Fig 5) are up-regulated; iii) *psbA* is up-regulated in expression for accelerated PSII turnover (Fig. 4); iv) To fight issues on protein level that come with mismetallation, such as misfolding or inhibited enzyme activity, protease and chaperones (e.g., FtsH1/2, GroEL1/L2/S, ClpB/B2, DnaJ/K1) are more efficiently transcribed (Fig. 5), as also indicated by KEGG map enrichment for chaperons and folding catalysts in the *Δmnx* mutant (Fig. 3D, Table S7). Overall, it was obvious that after Mn excess treatment transcript levels of proteins with rather stable pools, such as photosynthesis or antenna proteins, were down-regulated while in contrast transcripts related to ribosomes were enhanced (Fig. 3C, D). This finding indicates that cells rather invest in ribosomes (Fig. 3D, Fig. 7) likely to foster biosynthesis of proteins with higher turnover due to being either damaged by ROS (*e.g*., PsbA), mismetallated, or misfolded.

### One causative factor of Mn intoxication in Δ*mnx* is reduced energy metabolism

In contrast to the WT, the Δ*mnx* mutant was strongly negatively impacted by Mn in growth and pigmentation (Fig. 1). To investigate the nature/reason of Mn toxicity, the Δ*mnx* mutant is a reasonable study object since Mn efflux is hindered in this line due to the deletion of Mnx [10, 16]. An app. 3-fold enhanced intracellular Mn load could be demonstrated [10, 16]. Basically, the Δ*mnx* mutant mounts an adequate response on transcriptional level that corresponds with the WT response. Alterations that are exclusive to the mutant may indicate critical effects of Mn toxicity.

Besides transporters, Δ*mnx*-exclusive large changes on transcriptional level after Mn excess application were detected for the categories “photosynthesis” and “central carbon metabolism” (Fig. 3, Fig. 5, Fig. 7). Photosynthesis as the basis of the energy metabolism for oxygenic photosynthetic organisms is highly Fe-dependent: Chlorophyll biosynthesis is directly linked to heme biosynthesis and PSI together with the cytochrome-*b_6_f* complex requires a total of 12 Fe atoms, which are mainly used as cofactors or as Fe-S clusters [35]. Genes encoding proteins of the light reactions (PSI, PSII, phycobilisome, pigment biosynthesis, cytochrome *b_6_f* complex) were reduced in their transcript abundances in general under Mn excess conditions (Fig. 5; Table S7), as also indicated by the component loading of PC1 (Fig. 2B, Table S2, Table S3) and KEGG map enrichment (Fig. 3C). Reduction in pigment accumulation, that is chlorophyll, phycocyanin, and also carotenoids, was clearly detectable also on physiological level in the Δ*mnx* mutant after adding excess Mn (Fig. 1C, D, E) and is considered as a typical sign of Mn intoxication [61], as the reduction in PSI is [60]. Besides photosynthesis, our results furthermore showed that transcripts corresponding to the ATPase were significantly reduced in abundance (Fig. 3C; Fig. 5; Table S10). We postulate, light harvesting and photosynthetic electron transport become consequently impaired under Mn excess conditions, finally manifesting in reduced generation of ATP and reduction equivalents. The depletion in reduction equivalents goes hand in hand with impaired CO_2_ fixation due to reduced transcript levels of rib<u>u</u>lose-1,5-<u>bis</u>phosphate <u>c</u>arboxylase/<u>o</u>xygenase (Rubisco) subunits and enzymes of the CBB cycle (Fig. 3C; Fig. 5). Switching to heterotrophic life style is not assistant, since also genes encoding glycolytic enzymes are together with ATPase down-regulated (Fig. 3C; Fig. 5). As indicated by higher transcript levels of NDH-1 (Fig. 5), cells enhance cyclic electron transfer to match the reduced NADH+H^+^ consumption due to reduced CO_2_-fixation capacity via Rubisco and the CBB cycle but current cellular ATP need. In summary, our results from the RNA-seq analysis indicate a significant reduction of ATP generation and energy metabolism in general (Fig. 7).

### Oxidative stress is concurrently causative for Mn intoxication in Δ*mnx*

Another obvious differential feature of the transcriptional response in the Δ*mnx* mutant upon Mn treatment is the induction of the *sufBCDS* operon. The operon encodes for proteins of the SUF-system, which is central to the biogenesis of Fe-S cluster proteins in cyanobacteria and other bacteria (reviewed in [62]). Expression of the operon is under control of the transcription regulator SufR, which binds one [4Fe-4S]-cluster per subunit, forms homodimers as holoproteins, and acts as repressor upon binding. The Fe-S cluster functions as sensor for Fe availability and cellular redox status [63]. Oxidative stress causes damage to the SufR Fe-S clusters. As a result, the binding-affinity of SufR to the *sufBCDS*-promoter region is reduced and expression of the *sufBCDS* operon not any longer repressed [37, 63]. Accordingly, we interpret the enhanced expression of the operon as an indication for sensing of enhanced oxidative stress in Δ*mnx* as also noted by enhanced expression of ROS scavenging enzymes (Fig. 5, Fig. 7).

The enhanced ROS generation is possibly a consequence of the cytoplasmic Mn overload and concurrent mismetallation events, which impair photosynthetic electron transfer. Ample chaperones and folding catalysts are up in Δ*mnx* (Fig. 3D; Fig. 5) indicating tightened protein quality issues. However, while the WT was able to adjust cytosolic Mn homeostasis by Mnx-catalyzed Mn efflux, the mutant just tipped over the edge of tolerable cytoplasmic Mn concentration and the sum of protective mechanisms is not sufficient to alleviate Mn toxicity. The central importance of Mnx in maintaining cytoplasmic Mn homeostasis is highlighted by the observation that *Synechocystis* WT thrives even on 400 µM MnCl_2_ tested [10]. Without Mnx, the ultimate outcome of high cytoplasmic Mn load due to impaired Mn efflux is cell death (Fig. 7).

## Conclusions

The cyanobacterium *Synechocystis* tolerates treatments with high MnCl_2_ concentrations without negative effects on its performance. Respective transcriptional profiles indicate mismetallation of the canonical Fe-regulated transcription regulator Fur, enabling crossregulation of Mn- and Fe-responsive genes. Investment into ribosomes likely enable compensation of Mn-dependent mismetallation and protein damage. In case of impaired Mn efflux by Mnx, cytoplasmic Mn accumulation acts toxic by shutting-down central parts in energy metabolism covering both photosynthesis and respiration. Emerging ROS generation cannot be sufficiently compensated by protective measures making the effects of Mn intoxication in the Δ*mnx* mutant fatal. Our analyses thus reveal i) Mnx is not involved in sensing and transmission of cellular Mn status, but ii) Mnx is of absolute importance in balancing cytoplasmic Mn homeostasis.

## Acknowledgements

The authors thank the class (2022) of the master module “Methods and examples of functional genome research” at Bielefeld University for experimental support.

## Author contributions

M.R. and M.E. designed the experiments; M.R., P.V., and M.E., performed the experiments; P.V., K.N., A.B., and M.E. provided resources; M.R., S.Z., K. N., A.B. and M.E. analyzed the data; M.R. and M.E. wrote the manuscript with input from S.Z., K.N., and A.B.; all authors reviewed the final draft of the manuscript.

## REFERENCES

1. Chandler LE, Bartsevich V V., Pakrasi HB. Regulation of manganese uptake in *Synechocystis* 6803 by rfrA, a member of a novel family of proteins containing a repeated five-residues domain. Biochemistry 2003;42:5508–5514.

2. Sevilla F, López-Gorgé J, Gómez M, del Río LA. Manganese superoxide dismutase from a higher plant. Planta 1980;150:153–157.

3. Stengel A, Gügel IL, Hilger D, Rengstl B, Jung H, et al. Initial steps of photosystem II *de novo* assembly and preloading with manganese take place in biogenesis centers in *Synechocystis*. Plant Cell 2012;24:660–675.

4. Keren N, Kidd MJ, Penner-Hahn JE, Pakrasi HB. A light-dependent mechanism for massive accumulation of manganese in the photosynthetic bacterium *Synechocystis* sp. PCC 6803. Biochemistry 2002;41:15085–15092.

5. De Las Rivas J, Heredia P, Roman A. Oxygen-evolving extrinsic proteins (PsbO,P,Q,R): Bioinformatic and functional analysis. Biochim Biophys Acta Bioenerg 2007;1767:575–582.

6. Zhang B, Zhang C, Liu C, Jing Y, Wang Y, et al. Inner Envelope CHLOROPLAST MANGANESE TRANSPORTER 1 Supports Manganese Homeostasis and Phototrophic Growth in Arabidopsis. Mol Plant 2018;11:943–954.

7. Xiao Y, Huang G, You X, Zhu Q, Wang W, et al. Structural insights into cyanobacterial photosystem II intermediates associated with Psb28 and Tsl0063. Nat Plants 2021;7:1132–1142.

8. Eisenhut M. Manganese Homeostasis in Cyanobacteria. Plants 2020, *Vol 9*, *Page 18* 2019;9:18.

9. Duy D, Soll J, Philippar K. Solute channels of the outer membrane: From bacteria to chloroplasts. Biol Chem 2007;388:879–889.

10. Brandenburg F, Schoffman H, Kurz S, Krämer U, Keren N, et al. The *Synechocystis* manganese exporter Mnx is essential for manganese homeostasis in cyanobacteria. Plant Physiol 2017;173:1798–1810.

11. Waters LS. Bacterial manganese sensing and homeostasis. Current Opinion in Chemical Biology 2020;55:96–102.

12. Reis M, Brandenburg F, Knopp M, Flachbart S, Bräutigam A, et al. Hemi Manganese Exporters 1 and 2 enable manganese transport at the plasma membrane in cyanobacteria. bioRxiv 2023;2023.02.16.528846.

13. Yamaguchi K, Suzuki I, Yamamoto H, Lyukevich A, Bodrova I, et al. A two-component Mn^2+-^sensing system negatively regulates expression of the *mntCAB* operon in *Synechocystis*. Plant Cell 2002;14:2901–2913.

14. Zorina A, Sinetova MA, Kupriyanova E V., Mironov KS, Molkova I, et al. *Synechocystis* mutants defective in manganese uptake regulatory system, ManSR, are hypersensitive to strong light. Photosynth Res 2016;130:11–17.

15. Ogawa T, Bao DH, Katoh H, Shibata M, Pakrasi HB, et al. A two-component signal transduction pathway regulates manganese homeostasis in *Synechocystis* 6803, a photosynthetic organism. J Biol Chem 2002;277:28981–28986.

16. Gandini C, Schmidt SB, Husted S, Schneider A, Leister D. The transporter SynPAM71 is located in the plasma membrane and thylakoids, and mediates manganese tolerance in *Synechocystis* PCC6803. New Phytol 2017;215:256–268.

17. Salomon E, Keren N. Manganese Limitation Induces Changes in the Activity and in the Organization of Photosynthetic Complexes in the Cyanobacterium *Synechocystis* sp. Strain PCC 6803. Plant Physiol 2011;155:571–579.

18. Kaur G, Kumar V, Arora A, Tomar A, Ashish, et al. Affected energy metabolism under manganese stress governs cellular toxicity. Sci Rep 2017;7:1–11.

19. Irving J. P. H and W, Irving, H. and Williams JP. Order of stability of metal complexes. Nature 1948;162:746–747.

20. Foster AW, Osman D, Robinson NJ. Metal preferences and metallation. J Biol Chem 2014;289:28095–28103.

21. Rippka E, Deruelles J, Waterbury NB. Generic Assignments, Strain Histories and Properties of Pure Cultures of Cyanobacteria. J Gen Microbiol 1979;111:1–61.

22. Sigalat C, De Kouchkovsky Y. Fractionnement et caracterisation de l’appareil photosynthetique de l’algue bleue unicellulaire *Anacystis nidulans*. I. Obtention de fractions membranaires par “lyse osmotique” et analyse pigmentaire. Physiol Veg 1975;13:243–258

23. Robinson MD, McCarthy DJ, Smyth GK. edgeR: A Bioconductor package for differential expression analysis of digital gene expression data. Bioinformatics 2009;26:139–140.

24. Benjamini Y, Hochberg Y. Controlling the False Discovery Rate: A Practical and Powerful Approach to Multiple Testing. J R Stat Soc Series B Stat Methodol 1995;57:289–300.

25. Kanehisa M, Furumichi M, Tanabe M, Sato Y, Morishima K. KEGG: new perspectives on genomes, pathways, diseases and drugs. Nucleic Acids Res 2017;45:D353.

26. Cantalapiedra CP, Hern̗andez-Plaza A, Letunic I, Bork P, Huerta-Cepas J. eggNOG-mapper v2: Functional Annotation, Orthology Assignments, and Domain Prediction at the Metagenomic Scale. Mol Biol Evol 2021;38:5825–5829.

27. Moriya Y, Itoh M, Okuda S, Yoshizawa AC, Kanehisa M. KAAS: an automatic genome annotation and pathway reconstruction server. Nucleic Acids Res 2007;35:W182–W185

28. Fisher RA. On the Interpretation of χ 2 from Contingency Tables, and the Calculation of P. J R Stat Soc 1922;85:87.

29. Singh AK, McIntyre LM, Sherman LA. Microarray analysis of the genome-wide response to iron deficiency and iron reconstitution in the cyanobacterium *Synechocystis* sp. PCC 6803. Plant Physiol 2003;132:1825–1839.

30. Opiyo SO, Pardy RL, Moriyama H, Moriyama EN. Evolution of the Kdo2-lipid A biosynthesis in bacteria. BMC Evol Biol 2010;10:1–13.

31. Bartsevich VV, Pakrasi HB. Molecular identification of an ABC transporter complex for manganese: Analysis of a cyanobacterial mutant strain impaired in the photosynthetic oxygen evolution process. EMBO J 1995;14:1845–1853.

32. Nickelsen J, Rengstl B. Photosystem II assembly: From cyanobacteria to plants. Annu Rev of Plant Biol 2013;64:609–635.

33. Giner-Lamia J, López-Maury L, Reyes JC, Florencio FJ. The CopRS two-component system is responsible for resistance to copper in the cyanobacterium *Synechocystis* sp. PCC 6803. Plant Physiol 2012;159:1806–1818.

34. Hernández-Prieto MA, Schön V, Georg J, Barreira L, Varela J, et al. Iron deprivation in synechocystis: Inference of pathways, non-coding RNAs, and regulatory elements from comprehensive expression profiling. G3: Genes Genomes Genet 2012;2:1475–1495.

35. Kroh GE, Pilon M. Regulation of Iron Homeostasis and Use in Chloroplasts. Int J Mol Sci 2020, *Vol 21, Page 3395* 2020;21:3395.

36. Riediger M, Hernández-Prieto MA, Song K, Hess WR, Futschik ME. Genome-wide identification and characterization of Fur-binding sites in the cyanobacteria *Synechocystis* sp. PCC 6803 and PCC 6714. DNA Res 2021;28:dsab023.

37. Wang T, Shen G, Balasubramanian R, McIntosh L, Bryant DA, et al. The *sufR* Gene (sll0088 in S*ynechocystis* sp. Strain PCC 6803) Functions as a Repressor of the *sufBCDS* Operon in Iron-Sulfur Cluster Biogenesis in Cyanobacteria. J Bacteriol 2004;186:956.

38. Keren N, Aurora R, Pakrasi HB. Critical Roles of Bacterioferritins in Iron Storage and Proliferation of Cyanobacteria. Plant Physiol 2004;135:1666–1673.

39. Shcolnick S, Shaked Y, Keren N. A role for mrgA, a DPS family protein, in the internal transport of Fe in the cyanobacterium *Synechocystis* sp. PCC6803. Biochim Biophys Acta Bioenerg 2007;1767:814–819.

40. Hualing M. Cyanobacterial NDH-1 Complexes. Front Microbiol 2022;13:933160.

41. Mustila H, Muth-Pawlak D, Aro EM, Allahverdiyeva Y. Global proteomic response of unicellular cyanobacterium *Synechocystis* sp. PCC 6803 to fluctuating light upon CO_2_ step-down. Physiol Plant 2021;173:305–320.

42. Waters LS, Sandoval M, Storz G. The *Escherichia coli* MntR Miniregulon Includes Genes Encoding a Small Protein and an Efflux Pump Required for Manganese Homeostasis. J Bacteriol 2011;193:5887–5897.

43. Fisher CR, Wyckoff EE, Peng ED, Payne SM. Identification and characterization of a putative manganese export protein in *Vibrio cholerae*. J Bacteriol 2016;198:2810–2817.

44. Sharon S, Salomon E, Kranzler C, Lis H, Lehmann R, et al. The hierarchy of transition metal homeostasis: Iron controls manganese accumulation in a unicellular cyanobacterium. Biochim Biophys Acta Bioenerg 2014;1837:1990–1997.

45. Hernández-Prieto MA, Semeniuk TA, Giner-Lamia J, Futschik ME. The Transcriptional Landscape of the Photosynthetic Model Cyanobacterium *Synechocystis* sp. PCC6803. Sci Rep 2016 6:1 2016;6:1–15.

46. Salomon E, Keren N. Acclimation to environmentally relevant Mn concentrations rescues a cyanobacterium from the detrimental effects of iron limitation. Environ Microbiol 2015;17:2090–2098.

47. Lee JW, Helmann JD. Functional specialization within the fur family of metalloregulators. BioMetals 2007;20:485–499.

48. Ma Z, Faulkner MJ, Helmann JD. Origins of specificity and cross-talk in metal ion sensing by *Bacillus subtilis* Fur. Mol Microbiol 2012;86:1144–1155.

49. Jia A, Zheng Y, Chen H, Wang Q. Regulation and Functional Complexity of the Chlorophyll-Binding Protein IsiA. Front Microbiol 2021;12:774107.

50. Riediger M, Kadowaki T, Nagayama R, Georg J, Hihara Y, et al. Biocomputational Analyses and Experimental Validation Identify the Regulon Controlled by the Redox-Responsive Transcription Factor RpaB. iScience 2019;15:316–331.

51. Cheng D, He Q. PfsR Is a Key Regulator of Iron Homeostasis in *Synechocystis* PCC 6803. PLoS One 2014;9:e101743.

52. Peter Howard S, Herrmann C, Stratillo CW, Braun V. *In vivo* synthesis of the periplasmic domain of TonB inhibits transport through the FecA and FhuA iron siderophore transporters of *Escherichia coli*. J Bacteriol 2001;183:5885–5895.

53. Lau CKY, Krewulak KD, Vogel HJ. Bacterial ferrous iron transport: the Feo system. FEMS Microbiol Rev 2016;40:273–298.

54. Chenault SS, Earhart CF. Identification of hydrophobic proteins FepD and FepG of the *Escherichia coli* ferrienterobactin permease. J Gen Microbiol 1992;138:2167–2171.

55. Bosma EF, Rau MH, Van Gijtenbeek LA, Siedler S. Regulation and distinct physiological roles of manganese in bacteria. FEMS Microbiol Rev 2021;45:fuab028.

56. Martin JE, Waters LS, Storz G, Imlay JA. The *Escherichia coli* Small Protein MntS and Exporter MntP Optimize the Intracellular Concentration of Manganese. PLoS Genet 2015;11:e1004977

57. Bartsevich VV, Pakrasi HB. Membrane topology of MntB, the transmembrane protein component of an ABC transporter system for manganese in the cyanobacterium *Synechocystis* sp. strain PCC 6803. J Bacteriol 1999;181:3591–3593.

58. Bernal M, Casero D, Singh V, Wilson GT, Grande A, et al. Transcriptome Sequencing Identifies SPL7-Regulated Copper Acquisition Genes *FRO4/FRO5* and the Copper Dependence of Iron Homeostasis in Arabidopsis. Plant Cell 2012;24:738–761.

59. Imlay JA. The mismetallation of enzymes during oxidative stress. J Biol Chem 2014;289:28121–28128.

60. Millaleo R. Excess manganese differentially inhibits photosystem I versus II in Arabidopsis thaliana. J Exp Bot 2012;63:695–709.

61. Csatorday K, Gombos Z, Szalontai B. Mn^2+^ and Co^2+^ toxicity in chlorophyll biosynthesis. Proc Natl Acad Sci USA 1984;81:476–478.

62. Pérard J, Ollagnier de Choudens S. Iron–sulfur clusters biogenesis by the SUF machinery: close to the molecular mechanism understanding. J Biol Inorg Chem 2018;23:581–596.

63. Shen G, Balasubramanian R, Wang T, Wu Y, Hoffart LM, et al. SufR Coordinates Two [4Fe-4S]2+, 1+ Clusters and Functions as a Transcriptional Repressor of the *sufBCDS* Operon and an Autoregulator of *sufR* in Cyanobacteria. J Biol Chem 2007;282:31909–31919.

64. Mills LA, McCormick AJ, Lea-Smith DJ. Current knowledge and recent advances in understanding metabolism of the model cyanobacterium *Synechocystis* sp. PCC 6803. Biosci Rep 2020;40:20193325.

