## Supplementary material for "Study of excess manganese stress response highlights the central role of manganese exporter Mnx for holding manganese homeostasis in the cyanobacterium *Synechocystis sp.* PCC 6803": Fig. S1

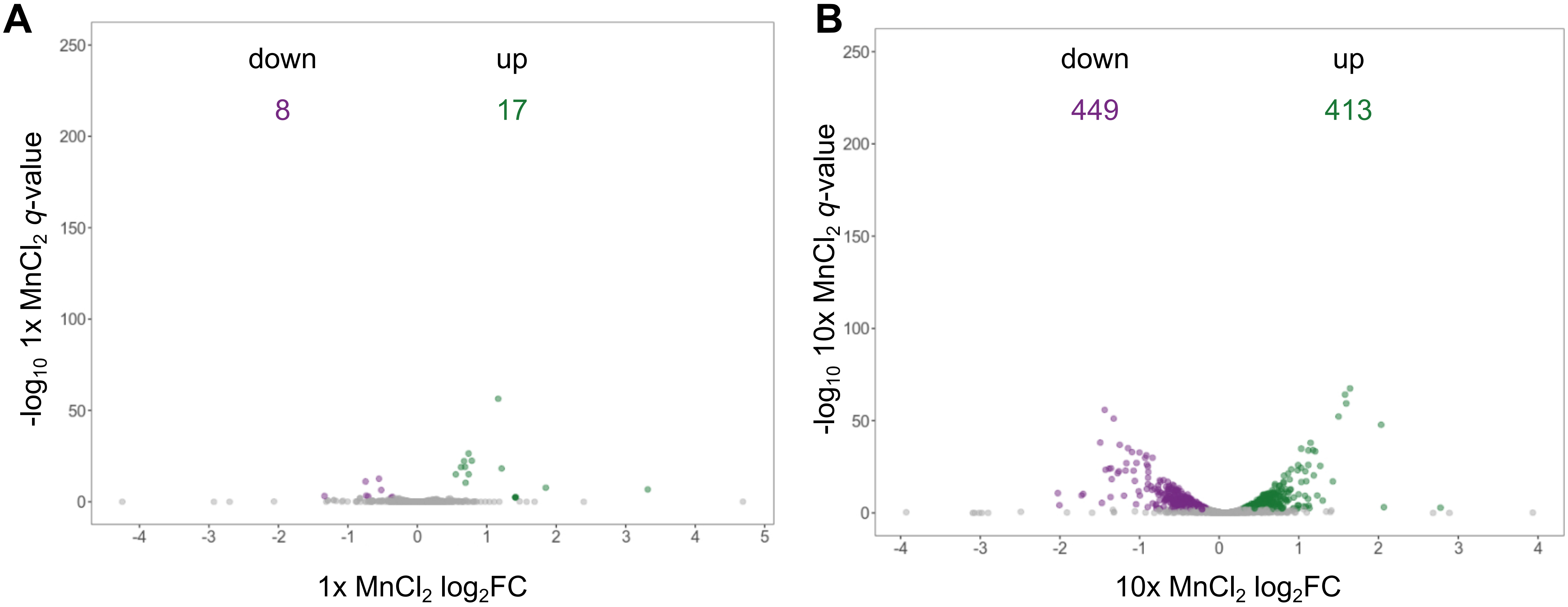


**Fig. S1:** Volcano plots of the global transcriptome responses under **(A)** control (1x $MnCl_2$) and **(B)** Mn excess (10x $MnCl_2$) conditions. Shown are $log_2$-fold changes ($log_2FC$) of Δ*mnx* mutant line *versus* WT under each condition. Differentially expressed genes ($q < 0.01$; edgeR, [24]) are plotted in green (up) or violet (down) respectively. The number of differentially expressed genes is given in violet for significantly downregulated and green for significantly upregulated genes.
